# Interactions between helicase and primase are crucial for DNA replication in the enteropathogen *Clostridium difficile*

**DOI:** 10.1101/071829

**Authors:** Erika van Eijk, Vasileios Paschalis, Matthew Green, Annemieke H. Friggen, Marilynn A. Larson, Keith Spriggs, Geoffrey S. Briggs, Panos Soultanas, Wiep Klaas Smits

**Affiliations:** Department of Medical Microbiology, Leiden University Medical Center, Leiden, the Netherlands; School of Chemistry, Center for Biomolecular Sciences, University of Nottingham, United Kingdom; Department of Microbiology and Pathology, University of Nebraska Medical Center, Omaha, NE 68198-6495; National Strategic Research Center, Omaha, NE 68198-4238; School of Pharmacy, University of Nottingham, United Kingdom.

**Author notes:** The authors wish it to be known that, in their opinion, the first 3 authors should be regarded as joint First Authors. To whom correspondence should be addressed. Department of Medical Microbiology, Leiden University Medical Center, PO Box 9600, 2300RC, Leiden, The Netherlands.

## Abstract

DNA replication is an essential and conserved process in all domains of life and may serve as a target for the development of new antimicrobials. However, such developments are hindered by a limited understanding of DNA replication in pathogenic micro-organisms. *Clostridium difficile* is the main cause of health-care associated diarrhea and its DNA replication machinery is virtually uncharacterized. We identified the replicative helicase (CD3657), the helicase loader ATPase (CD3654) and primase (CD1454) of *C. difficile* and reconstitute helicase and primase activity *in vitro*. We demonstrate a direct and ATP-dependent interaction between the helicase loader and the helicase. We find that helicase activity is dependent on the presence of primase *in vitro*. The trinucleotide specificity of primase, which we find to be determined by a single lysine residue, is similar to extreme thermophile *Aquifex aeolicus* but the presence of helicase allows more efficient *de novo* synthesis of RNA primers from non-preferred trinucleotides. Thus, helicase-primase interactions constitute a crucial aspect of DNA replication initiation in *C. difficile* and helicase loading and activation in this organism differs critically from that of the Gram-positive model organism *Bacillus subtilis*.

## INTRODUCTION

Extensive research, primarily on the model organisms *Escherichia coli* (Gram-negative) and *Bacillus subtilis* (Gram-positive), has shown that many different proteins are involved in DNA replication. Although the overall mechanism of replication is highly conserved in all domains of life, it is perhaps not surprising that details of the molecular mechanisms can vary substantially as these prokaryotes diverged more than 3 billion years ago (1).

One of the best characterized distinctions between *B. subtilis* and *E. coli* is the mechanism of loading the replicative helicase at the origin of replication (*oriC*), an essential step in the DNA replication process of bacteria (2;3). In the Enterobacteria, Firmicutes and Aquificae this event is facilitated by a specific loader protein, which is not conserved in bacteria outside these phyla (4). However, the strategy of helicase loading among bacteria that do encode a loader protein differs (5;6).

For historical reasons, the nomenclature for the replication proteins differs between bacterial species (e.g. *E. coli* helicase; DnaB and *B. subtilis* helicase; DnaC). For clarity, protein names hereafter will be used in conjunction with species and either written in full or abbreviated (e.g. Ec, Bs). The *E. coli* helicase (EcDnaB) is loaded by a single loader protein (EcDnaC) *in vivo* (7-9), whereas loading of the *B. subtilis* helicase (BsDnaC) requires three accessory proteins (BsDnaD, BsDnaB and BsDnaI) *in vivo* (10-13) in addition to the replication initiator DnaA that is required in both organisms. One possible explanation for the requirement of multiple proteins in *B. subtilis* may lie in the fact that *E. coli* and *B. subtilis* employ different mechanisms to deliver the replicative helicase onto the DNA (5;6). Alternatively, it may reflect different *oriC* architectures, requiring different mechanisms of origin remodelling (14).

Replicative helicases form hexameric rings and require single-stranded DNA (ssDNA) to be threaded through the central channel of the protein to to unwinding the DNA duplex (6;15). To accomplish this, it is thought that either the preformed ring is physically opened (ring-breaker) or that the ring is assembled from monomers at *oriC* (ring-maker) (5).

In *E. coli*, preformed hexamers of the helicase protein are capable of self-loading onto ssDNA. They display *in vitro* translocation and unwinding activities, which are highly induced in the presence of the loader protein (8). This in contrast to *B. subtilis*, where pre-assembled hexameric helicase is inactive, irrespective of the presence of the loader protein. *In vitro*, *B. subtilis* helicase activity is only observed when the helicase protein is monomeric and the loader protein is present (16). Thus, helicase loading in *E. coli* is an example of the ring breaker mechanism, whereas the situation in *B. subtilis* exemplifies a ring maker mechanism (5).

*B. subtilis* helicase loading *in vivo* is a hierarchical process (11-13). Initially, the double-stranded DNA (dsDNA) at *oriC* is melted into ssDNA by the initiation protein DnaA, thereby creating a substrate for primosome assembly. The BsDnaD and BsDnaB co-loader proteins, which are structural homologs (PFAM DnaB_2), associate sequentially with the replication origin(11) and possibly contribute to origin remodelling. This ultimately enables the ATPase loader protein to load the helicase (11-14;17-21).

The replication initiation protein (BsDnaA) and helicase loader protein (BsDnaI) belong to the AAA+ (ATPases associated with various cellular activities) family of ATPases (22-25). These AAA+ enzymes are, in their turn, part of the additional strand catalytic glutamate (ASCE) family (14;25-29). The BsDnaI loader protein consists of a C-terminal AAA+ domain that is necessary for nucleotide and ssDNA binding, and an N-terminal helicase-interacting domain (16). The loader interacts with the helicase regulating its activity and is therefore pivotal in replisome assembly (3;30;31).

The BsDnaC helicase is also an ASCE protein, and belongs to RecA-type helicase Superfamily 4 (SF4), that is involved in DNA replication (27;32;33). The SF4 superfamily of helicases is characterized by five sequence motifs; H1, H1a, H2, H3, and H4 (27;34;35). Motifs H1 and H2 are equivalent to the ATP-coordinating Walker A and B motifs found in many other ATPases (27).

The multi-protein primosome of *B. subtilis* consists not only of the helicase loader protein and helicase, but also of primase (36;37). Primase is pivotal for the initiation of DNA-synthesis at the replication origin and remains of utmost importance during the DNA-replication process in restarting stalled replication forks as well as *de novo* priming of Okazaki fragments for lagging strand synthesis (38). Prokaryotic primases have a three-domain structure consisting of an N-terminal zinc-binding domain, a central RNA polymerase domain that catalyzes the polymerization of ribonucleotides and a C-terminal domain that either is responsible for the interaction with helicase (helicase interaction domain) or has helicase activity itself (39). The latter region, also known as P16 reflecting the approximate mass (16kDa), is variable in prokaryotes and seems to be crucial for the direct interaction with and activation of the helicase (40-43). Interestingly, the P16 domain is structurally and functionally homologous to the N-terminal domain of the replicative helicase to which it binds (42;44;45). Depending on the bacterial species, the interaction between helicase and primase can be either transient (*E. coli*) or stable (*Geobacillus stearothermophilus*) (40;46).

Primase and helicase affect each other’s activities (40;43;47-49). Helicase affects primase by modulating initiation specificity, stimulating primer synthesis, reducing the length of primers synthesized and increasing its affinity for single stranded DNA (39;43;47-54). Although increased activity of primase in the presence of helicase was shown for Firmicutes such as *Staphylococcus aureus* and *B. subtilis*, primer length was minimally or not altered in either organism (49;55;56). Conversely, ATPase and helicase activities of helicase are stimulated by primase due to stabilization of the helicase hexamer by this interaction (40;43;57-59). In *Helicobacter pylori*, it was shown that the interaction of primase with helicase resulted in dissociation of the double hexamer of helicase thereby increasing ATP hydrolysis, DNA binding and unwinding (15).

Although no interaction of primase with loader protein has been demonstrated to date, it has been suggested that primase affects the interaction between helicase and the loader protein in *E. coli* (60). Dissociation of loader protein from the C-terminal region of helicase in this organism is thought to be induced by primer synthesis and conformational changes resulting from primase-helicase interaction (60). Similar observations were made in *B. subtilis*, where the loader protein was found to dissociate from a complex when primase and polymerase bind to helicase in gel-filtration experiments (56). However, a ternary complex comprising of helicase, loader, and the helicase binding domain of primase in *G. stearothermophilus* is capable of loading (61). These observations indicate that primosome-formation, like helicase loading, may be species-specific as well (58;60;61).

We are interested in DNA replication in the Gram positive enteropathogen *C. difficile,* the most common causative agent of antibiotic associated diarrhea (62;63). Like *B. subtilis*, *C. difficile* belongs to the low-GC content Firmicutes. This strictly anaerobic bacterium can cause potentially fatal intestinal inflammation (colitis), is resistant to multiple antibiotics, and is capable of forming highly resistant endospores (62;63). However, the *C. difficile* DNA replication machinery is virtually uncharacterized.

Herein we address this gap by identifying and characterizing the *C. difficile* replicative helicase (CD3657), helicase loader (CD3654) and primase (CD1454), and reconstituting helicase and primase activity *in vitro*. Our results indicate that the interactions between *C. difficile* helicase and primase are crucial for the functionality of both proteins, that a single residue can strikingly change trinucleotide specificity and that helicase loading and activation, as well as template specifity, in *C. difficile* differs critically from the *B. subtilis* model.

## MATERIAL AND METHODS

Supplementary Methods are available with further details.

### Plasmid construction and site directed mutagenesis

All oligonucleotides and plasmids constructed for this study are listed in **Appendix Tables S1 and S2**.

To construct the CD3654, CD3657 and CD1454 expression plasmids, the open reading frames were cloned into suitable pET vectors (Novagen) to yield pEVE24, pEVE87 and pEVE7, respectively.

Construction of the plasmids for the bacterial two hybrid system was performed with Gateway cloning technology (Invitrogen). To construct the CD3654 and CD3657 entry plasmids, the CD3654 and CD3657 open reading frames were introduced into donor vector pDonR™201 Bacterial two hybrid constructs were made by sub-cloning the genes of interest from the entry plasmids into the destination plasmids pKEK1286 (Zif fusion plasmid) or pKEK1287 (ω fusion plasmid) in an LR reaction (68), to yield pEVE118, pEVE120, pEVE122, pEVE123, pEVE124 and pEVE125.

Mutations in plasmids carrying CD3654, CD3654 and CD1454 mutants were constructed according to the QuikChange protocol (Stratagene). Primers were generated with Primer X, a web-based tool for automatic design of mutagenic primers for site-directed mutagenesis.

All constructs were verified by DNA sequencing.

### Purification of proteins

Overexpression of all proteins was carried out in *E. coli* BL21 (DE3) from the pET-based vectors pEVE12, pEVE90, pEVE92, pEVE24, pEVE59, pEVE60, pEVE203, pEVE7, and pEVE201 (**Appendix Table 2**).

Hexameric CD3657 was purified on a Q sepharose column, a heparin sepharose column and a Hiload 26/60 Superdex 200 gel filtration column. Monomers were recovered after the portion-wise addition of guanidinium chloride solution (8M), followed by separation of monomers and multimers on a Hiload 26/60 Superdex 200 column and and stored in TED50G20 buffer (Tris pH 7.5 50 mM, EDTA 1 mM, DTT 1 mM, NaCl 50 mM, 20% v/v glycerol) at -80°C after extensive dialysis.

CD3654 was purified on from the flow through of a Q sepharose and 5 mL SP sepharose columns connected in series, followed by a heparin sepharose and a Hiload 26/60 Superdex 200 gel filtration column. Purified CD3654 was stored in TED50G10 buffer (Tris pH 7.5 50 mM, EDTA 1 mM, DTT 1 mM, NaCl 50 mM, 10% v/v glycerol) at -80°C.

CD1454 was purified on HiTrap Q HP column in series with a 5ml heparin sepharose column, followed by a MonoS column and a Hiload 26/60 Superdex 200 column. Protein was stored in TED100G buffer (50 mM Tris pH7.5, 1 mM EDTA, 1 mM DTT, 100 mM NaCl, 10% v/v glycerol) at -80 °C.

Proteins were quantified by UV spectrophotometry and stored at -80°C. Protein purity (>95%) was estimated by SDS-PAGE electrophoresis and concentration was determined spectrophotometrically using extinction coefficients calculated by the ExPASy ProtParam tool (http://web.expasy.org/protparam).

#### Gel-filtration experiments

Purified proteins were studied in the presence and absence of 1 mM ATP on a Hiload 10/300 GL Superdex 200 analytical grade size exclusion column at a flow rate of 0.5 mL/min. The elution profiles from each experiment were monitored at 280nm and plotted as a function of the elution volume. To assess interactions between CD3657 and CD3654, purified proteins were mixed in a 1:1 stoichiometry. Samples from fractions were analysed by SDS-PAGE and Coomassie Blue staining to verify the identity of the proteins.

#### Bacterial two-hybrid assays

To determine (self-) interactions, pKEK1286- and pKEK1287-derived constructs were subsequently transformed in to the *E. coli* reporter strain KDZif1ΔZ (68;98) and assayed for β-galactosidase activity. Experiments were performed in triplicate.

#### Helicase assays

Helicase activity was assayed by monitoring (and quantifying) the displacement of the radiolabelled (γ^32^P-ATP) 105-mer oligonucleotide oVP-1 (partially) annealed to the single stranded circular DNA m13mp18 (ssM13; Affymetrix) essentially as previously described (56). All reactions, containing 0.658 nM radiolabelled DNA substrate, were initiated by the addition of 2.5 mM ATP and carried out at 37°C in buffer containing 20 mM HEPES-NaOH (pH 7.5), 50 mM NaCl,, 10 mM MgCl_2_ and 1 mM DTT for various lengths of time. The reactions were terminated by adding 5x SDS-STOP buffer (100 mM Tris pH8.0, 200 mM EDTA, 2.5% w/v SDS, 50% v/v glycerol, 0.15% (w/v) bromophenol blue). Proteins were added sequentially, with a 5 minute preincubation after each addition. The gel was dried, scanned and analyzed using a molecular imager and associated software (Biorad). Experiments were carried out in triplicate, and data analysis was performed using Prism 6 (GraphPad Software).

##### RNA primer synthesis assay and thermally denaturing HPLC analysis

RNA priming assays and denaturing HPLC analyses were conducted as was previously described for other mesophilic bacterial primases (73). The purified oligonucleotide templates used in this study were synthesized by Integrated DNA Technologies, Inc. (Coralville, IA). All RNA primer synthesis reactions were carried out in 50 µL nuclease-free water containing 50 mM HEPES pH 7.5, 100 mM potassium glutamate, 10 mM DTT, 2 μM ssDNA template, 30 mM magnesium acetate, and 1.2 mM of each NTP, unless otherwise specified. For the denaturing HPLC analyses, a gradient 0–8.8% v/v acetonitrile over 16 min was used to obtain optimal separation of primer products and ssDNA templates on a WAVE HPLC Nucleic Acid Fragment Analysis System equipped with a DNA Sep HPLC column (Transgenomic; Omaha, NE). The moles of RNA primers synthesized were quantified as previously described (55).

## RESULTS

### In silico *identification of putative replication initiation proteins*

In *B. subtilis*, replication initiation requires the coordinated action of multiple proteins, DnaA, DnaD, DnaB and DnaI, *in vivo* (11;14). BLASTP queries of the genome of *C. difficile* 630 (Genbank AM180355.1) using the amino acid sequence of these proteins from *Bacillus subtilis* subsp. *subtilis* strain 168 (GenBank NC0989.1) allowed the identification of homologs of most, but not all, proteins that were found to be essential for replication initiation in *B. subtilis*.

Homologues of the initiation protein DnaA (CD0001; e-value 0.0) and the putative helicase (CD3657; e-value 7e-713) were identified with high confidence, sharing respectively 62 and 52 percent identity with their *B. subtilis* counterparts across the full length of the protein. Interestingly, no homolog of BsDnaB was found using this strategy. However, the genome of *C. difficile* does harbour two homologs of BsDnaD (CD2943: E-value=2e-05, identity =29%, query coverage= 32%; CD3653: E-value=4e-05, identity 29%, query coverage 47%). As BsDnaB and BsDnaD are strictly required for replication initiation in *B. subtilis* and are structurally related (17), we further examined the *C. difficile* homologs of BsDnaD. BsDnaD is composed of two domains; DDBH1 and DDBH2 (PFAM DnaB_2), whereas BsDnaB has a DDBH1-DDBH2-DDBH2 structure (17). The DnaD-like proteins CD3653 and CD2943 of *C. difficile* both consist of three domains (DDBH1-DDBH2-DDBH2) and therefore resemble BsDnaB in domain structure (Figure 1A). CD2943 is annotated as a putative phage replication protein and is located in the ~50kb prophage 2 (64). Also in *Listeria monocytogenes*, *S. aureus* and *Lactobacillus plantarum* DDBH2-containing phage genes have been identified (17;65). Considering the chromosomal location, we consider a role for CD2943 in chromosomal DNA replication unlikely, but a role for CD3653 plausible.

**Figure 1.**
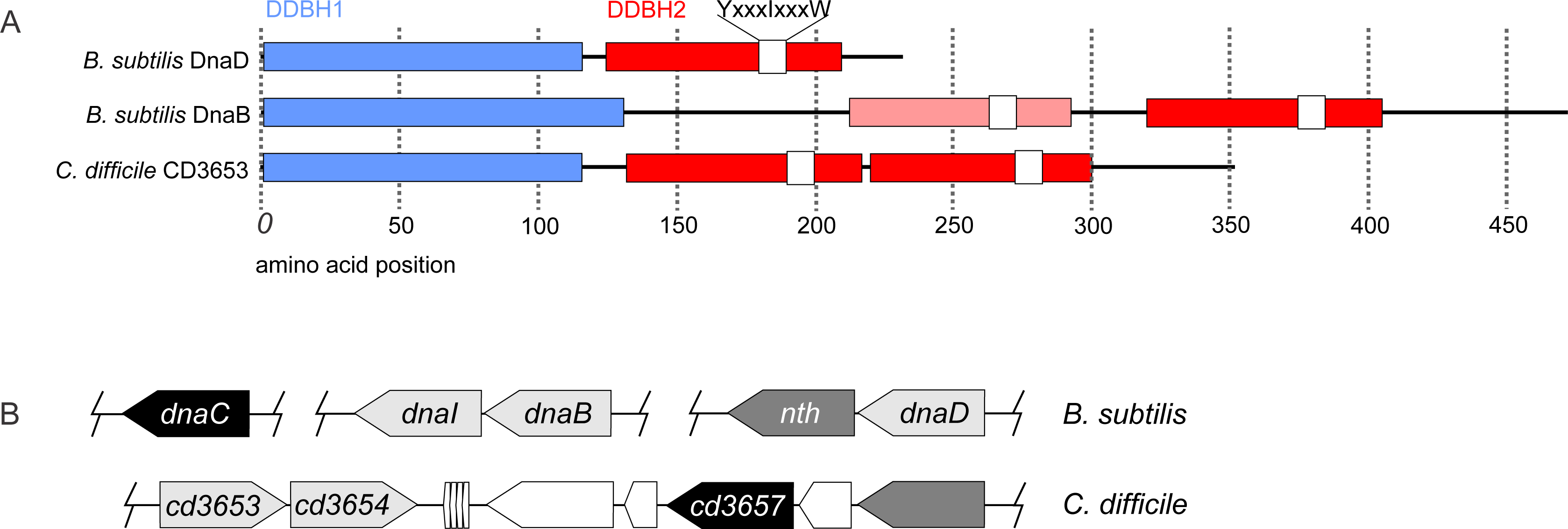
*In silico* analysis of putative replication initiation proteins of *C. difficile*. **A**. Domain structure of BsDnaD, BsDnaD and CD3653. Domain nomenclature according to Marston et al. (17). Note that DDBH2 corresponds to PFAM DnaB_2. **B**. Chromosomal organization of the *dnaBI* genomic region of *B. subtilis* and the CD3653-CD3654 genomic region of *C. difficile*.

A similar argument can be made for the putative helicase loader. Two homologues of the BsDnaI protein were identified in *C. difficile* (CD3654: E value=8e-10, identity 26%, query coverage=46%; CD0410: E-value=1e-18, identity=31%, query coverage=51%). The sequence homology of putative loader proteins with their counterpart in *B. subtilis* is mainly confined to the C-terminal AAA+ domain that contain the Walker A and B motifs (58). CD0410 is located on the conjugative transposon CTn*2* (66) and therefore not part of the core genome of *C. difficile*. The *dnaD*-like gene CD3653 is located adjacent to the putative loader (CD3654), in the same genomic region as the replicative helicase (CD3657) (Figure 1B). Of note, the *dnaB* gene of *B. subtilis* is located next to the gene helicase loader *dnaI* (67), suggesting a functional relationship between the loader ATPase and a DnaB_2 family protein. Indeed, it has been suggested that BsDnaB is a co-loader of the BsDnaC helicase (13). Taken together, our analyses strongly suggest that CD3654 is the cognate loader protein for the *C. difficile* replicative helicase CD3657.

### Helicase can form hexamers at high concentration

A distinguishing feature of the different modes of helicase loading (ring-maker vs. ring-breaker) is the multimeric state of helicase at dilute concentrations of protein (5). Therefore, we purified recombinant *C. difficile* helicase protein and determined its multimeric state using analytical gel filtration. At concentrations below 5 µM we observed predominantly monomeric protein, with a fraction of the protein forming low molecular weight complexes (likely dimers or trimers) while at 10 µM and above, the helicase formed hexamers (**Figure S1**). Thus, at physiological (nM) concentrations the *C. difficile* helicase is predominantly monomeric, suggesting it is of the ring maker type, like *B. subtilis*. Multimerization at high concentrations of protein was independent of the presence of ATP (unpublished observations).

To confirm the gel filtration data, we investigated the self-interaction of helicase in a bacterial two-hybrid system (68). This system detects interactions between a protein fused to Zif (Zinc-finger DNA binding domain) and a protein fused to the RNA polymerase ω subunit. Interaction between proteins of interest facilitates transcriptional activation of a Zif-dependent *lacZ* reporter gene in a dedicated reporter strain (68). In order to quantify the interaction, *E. coli* cells containing the plasmids encoding the fusion proteins were permeabilized and assayed for β-galactosidase activity. We found that in the reporter strain transformed with plasmids harboring both fusion proteins, β-galactosidase activity was ~3 fold higher than for the reporter strains harboring the individual plasmids, indicating a clear self-interaction for the *C. difficile* helicase protein (Figure 2A).

**Figure 2.**
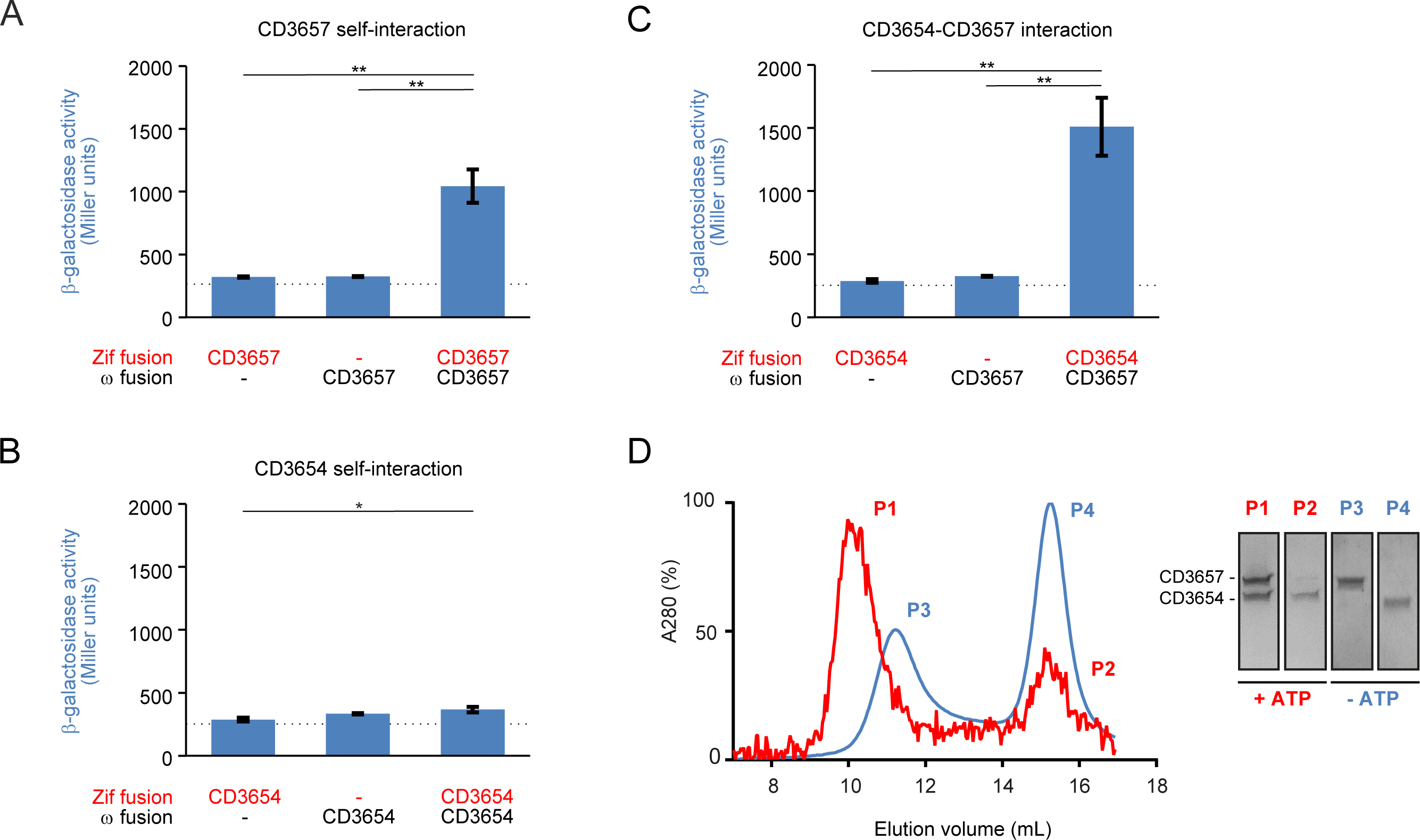
CD3654 and CD3657 interact. **A.** Bacterial two hybrid analysis of CD3657 self-interaction. **B**. Bacterial two hybrid analysis of CD3654 self-interaction. **C**. Bacterial two hybrid analysis of the CD3657-CD3654 interaction. Bar graphs in **A-C** indicate average values and error-bars indicate standard deviation of the measurements (n=3). Significance was determined using the Student’s t-test (* p< 0.05, ** p<0.001). **D.** CD3657 and CD3654 interact in an ATP-dependent manner. Analytical gel filtration was performed on a HiLoad 10/300 GL Superdex analytical grade size exclusion column with 2.21 μM of CD3657 and CD3654 in the presence (red) and absence (blue) of 1 mM ATP.

Similar experiments were carried out with the putative helicase loader (CD3654). Analytical gel filtration using purified loader protein (CD3654) showed that it was monomeric at all concentrations tested (unpublished observations). Consistent with this observation, no self-interaction of CD3654 was found in the bacterial two-hybrid system (Figure 2B).

We conclude that helicase can form homomultimeric assemblies, whereas the putative loader is monomeric under the conditions tested.

### Helicase and the putative helicase loader interact in an ATP-dependent manner

If CD3654 is a legitimate loader for the *C. difficile* helicase (CD3657), we expect that the proteins interact *in vivo* and *in vitro*. To determine if this is the case, we performed bacterial two-hybrid and analytical gel filtration experiments. First, we tested if an interaction between the helicase CD3657 and the putative loader CD3654 could be demonstrated in the bacterial two-hybrid system. CD3657 was fused to the RNA polymerase ω subunit and CD3654 was fused to Zif (Karna 2010). The β-galactosidase activity in the reporter strain containing both plasmids was ~5 -fold increased (p<0.001) compared to the reporter strains containing the individual plasmids (background) (Figure 2C). The high β-galactosidase activity implies substantial interactions between the *C. difficile* helicase and putative helicase loader. Similar results were obtained when CD3657 was fused to Zif and CD3654 to the RNA polymerase ω subunit (unpublished observations). This indicates that the combination of protein and fusion domain does not influence the results of this assay.

To exclude the possibility for false negative or false positive results as a result of the two-hybrid system, we additionally performed analytical gel filtration experiments using purified non-tagged CD3657 and CD3654 proteins. In these experiments, *C. difficile* helicase and loader were combined in equimolar concentrations (2.21 µM) in the presence and absence of ATP (1 mM). In the presence of ATP, the elution profile showed a major high molecular weight (MW) peak (~10 mL; ~500 kDa, P1) and a minor low MW peak (~15 mL; ~40 kDa, P2) (Figure 2D). In combination with a visual inspection of the fractions collected from both peaks on an Coomassie-stained SDS-PAGE gel, we believe that the major peak can be attributed to a dodecameric complex consisting of six CD3657 monomers and six CD3654 monomers (theoretical MW 525 kDa), whilst the minor peak corresponds to predominantly free monomeric CD3654 (theoretical MW 38 kDa). Similar results were obtained when high concentrations of proteins (~10 uM) were used (**Figure S2**), suggesting that pre-formed hexameric helicase retains the ability to interact with the CD3654 protein at the same stoichiometry.

The elution profile of the same concentration of proteins in the absence of ATP showed a completely different picture (Figure 2D). A minor peak was observed at ~11 mL (~300 kDa, P3) and a major second peak eluted at ~15 mL (~40 kDa, P4). Molecular weight estimates, in combination with an evaluation of peak fractions on an SDS-PAGE gel, indicated that the first peak most likely corresponds to a complex of six monomers of CD3657 (theoretical MW 297 kDa), whilst the second peak corresponds to monomeric CD3654 protein.

Together, the data shows that the CD3657 helicase and the loader CD3654 can form a complex in an ATP-dependent manner.

### Mutation of the helicase Walker A motif abrogates protein-protein interactions

To address the question which of the proteins (or both) requires ATP to promote the formation of a CD3657-CD3654 complex, mutants in the Walker A motif of both proteins were created. The Walker A motif (GXXXXGK[T/S]) directly and indirectly interacts with ATP, and is the principal ATP binding motif of P-loop ATPases (25). The motif is highly conserved in both helicase and helicase loader proteins (6). The lysine residue (K) forms a direct interaction with the negatively charged nucleotide β or γ phosphate group and mutation of this residue is known to abrogate nucleotide binding and lead to inactivation of P-loop ATPases. The threonine (T) residue in the Walker A motif either directly or indirectly coordinates a Mg^2+^ within the ATP binding site, which in turn coordinates the phosphate groups of ATP. In the *B. stearothermophilus* replicative helicase, mutation of the threonine residue results in a protein that lacks ATPase and unwinding activities (58). Based on this knowledge, the equivalent residues were identified in the *C. difficile* helicase protein. Using site-directed mutagenesis we generated mutant helicase proteins in which the lysine at position 214 was changed into an arginine (K214R) and the threonine at position 215 was changed into an alanine (T215A).

To determine whether these CD3657 proteins showed altered protein-protein interactions, we performed bacterial two hybrid experiments. We fused the CD3657 protein to the ω-subunit and CD3654 or CD3657 proteins to the Zif subunit and evaluated their ability to drive the expression of a transcriptional reporter. For both the CD3657 K214R and CD3657 T215A mutants, interaction with the wild type CD3654 protein were severely reduced or completely lost (Figure 3A-B). This prompted us to investigate if the mutant helicases still had the capacity to form homomultimeric assemblies, as it has been shown in other bacteria that oligomerization is an important step in the mechanism of action of DNA helicases (69). We found that the CD3657 K214R also demonstrated reduced self-interactions, whereas the CD3657 T215A mutant had completely lost the ability to self-interact (Figure 3C-D). We conclude that the ability of the CD3657 helicase to coordinate ATP correlates with its ability to interact with the putative loader CD3654 and the ability to self-interact. Since the most dramatic effect was observed for CD3657 T215A, we focused our further experiments on this particular mutant.

**Figure 3.**
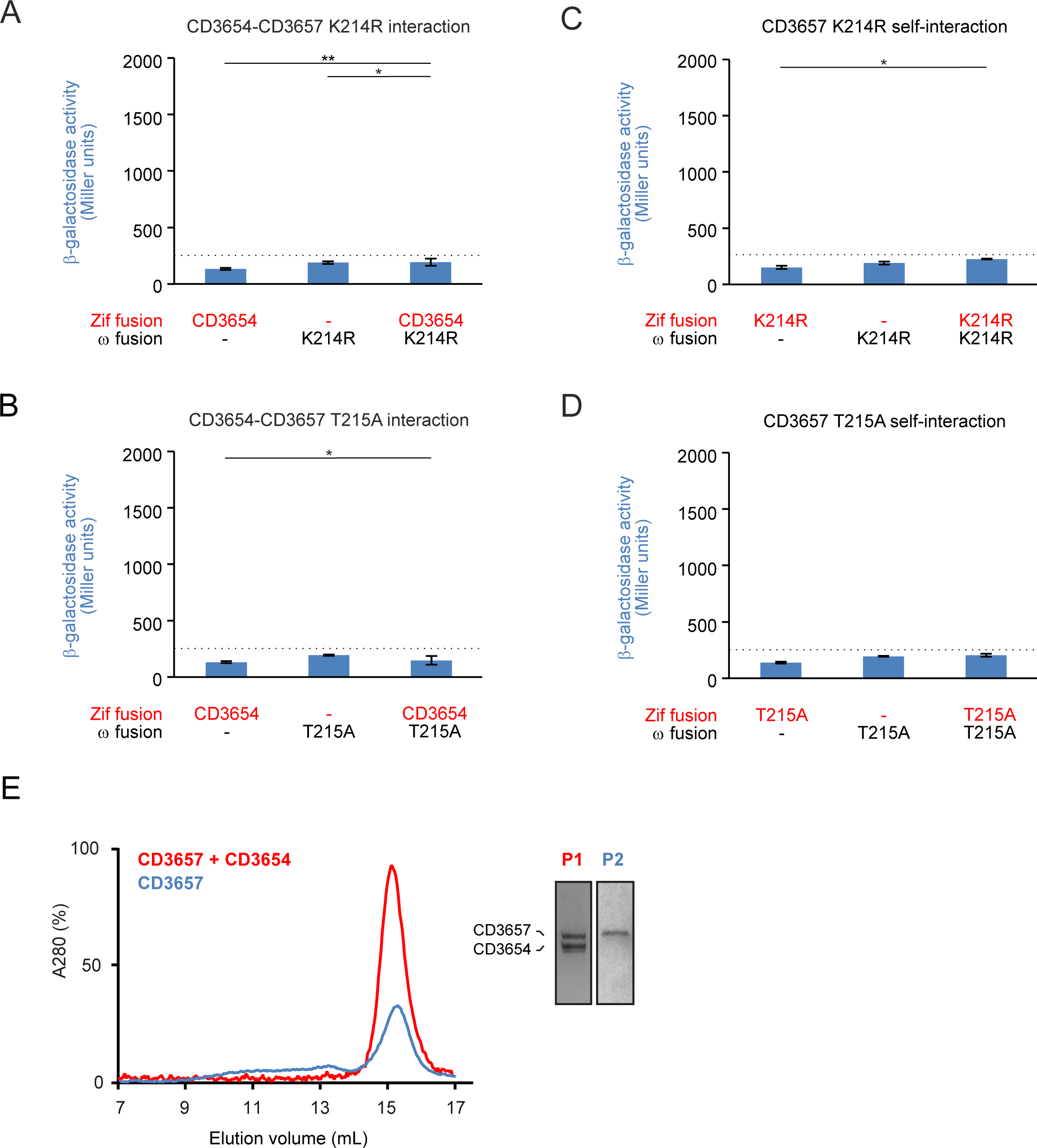
Walker A mutants of CD3657 are defective in protein-protein interactions. **A-B**. Walker A mutants of CD3657 (**A**. K214R; **B**. T215A) show no or severely reduced interactions with the putative loader protein CD3654 in a bacterial two hybrid assay. **C-D**. Walker A mutants of CD3657 (**C**. K214R; **D**. T215A) no longer self-interact in a bacterial two hybrid assay. Bar graphs in **A-D** indicate average values and error-bars indicate standard deviation of the measurements (n=3). Significance was determined using the Student’s t-test (* p< 0.05, ** p<0.001). **E**. CD3657 T215A is defective in protein-protein interactions. Analytical gel filtration was performed on a HiLoad 10/300 GL Superdex 200 analytical grade size exclusion column with 2.43 μM of CD3657 T215A in the presence (red) and absence (blue) of 2.43 μM CD3654.

To confirm the findings from the bacterial two hybrid experiments, we purified CD3657 T215A protein and subjected it to size exclusion chromatography (Figure 3E). *C. difficile* helicase T215A mutant (2.43 µM) was incubated in the presence of ATP (1 mM) and loaded onto a size exclusion column. A major peak was observed at ~15 mL (~40 kDa), likely corresponding to monomeric CD3657 T215A (theoretical MW 49 kDa). We did not observe any high MW complexes under these conditions, in contrast to the wild type CD3657 protein (Figure 2D). Next, we combined the CD3657 T215A mutant and the wild type CD3654 protein (both 2.43 µM) in the presence of ATP (1 mM) (Figure 3E). A single peak was observed at ~15 mL (~40 kDa). Analysis of the peak fractions on SDS-PAGE demonstrated that the peak contained both CD3657 T215A and CD3654 protein and thus corresponds to monomeric forms of both proteins (theoretical MW 49 and 38 kDa, respectively). Also in these experiments no high MW complexes were found, in contrast to the wild type CD3657 and CD3654 proteins (Figure 2D).

We also generated constructs with Walker A mutations in CD3654 (K198R, T199A) and tested these in the bacterial two hybrid assay and in size exclusion chromatography. The mutant loader proteins retained the ability to interact with the wild-type helicase protein (**Figure S3**). Self-interaction was not observed for the mutant loader proteins, concordant with the results obtained with the wild-type loader protein (unpublished observations).

Overall, our data show that the Walker A mutant CD3657 T215A can no longer self-interact and has lost the capacity to interact with the putative loader protein CD3654. We conclude that the ATP requirement for the interaction between the two proteins is most likely the result of ATP binding to CD3657.

### *Helicase loading of* C. difficile *differs from* B. subtilis

So far, our data show that the replicative helicase of *C. difficile*, CD3657, interacts in an ATP dependent manner with the helicase loader protein CD3654 and that loading likely occurs via a ring-maker mechanism, like for *B. subtilis*. In *B. subtilis*, stimulation of activity of the (monomeric) helicase protein by the loader protein was clearly shown using an *in vitro* helicase activity assay (16;56). Therefore, we set out to investigate if the activity of *C. difficile* helicase could be reconstituted in the presence of the CD3654 protein in a similar experiment. DNA unwinding helicase activity was assayed by monitoring and quantifying the displacement of a radiolabelled oligonucleotide annealed to single stranded circular M13mp18 DNA. To enable loading of helicase, the 5’ end of the oligonucleotide contained a poly(dCA) tail that produces a forked substrate upon annealing of the complementary region to ssM13. Wild type CD3657 and CD3654 proteins were mixed in equimolar concentrations in the presence of ATP and reaction buffer and displacement of the radiolabeled oligonucleotide was monitored over time. In contrast to the *B. subtilis* proteins, helicase activity was not observed during this time course (**Figure S4**). This suggests that another factor is required for *in vitro* loading and/or activation of the *C. difficile* helicase.

### *Identification of the* C. difficile *primase*

Primase has been shown to interact with helicase and stimulate its activity in a variety of organisms (15;56;60). Therefore, we wanted to investigate if *C. difficile* helicase could be activated by primase.

A BLASTP query (http://blast.ncbi.nlm.nih.gov/) of the *C. difficile* genome (GenBank AM180355.1) allowed the identification of a protein homologous to *B. subtilis* primase BsDnaG (GenBank NC_009089.1). This protein, CD1454, shared 31 percent identity with its *B. subtilis* counterpart across the full length of the protein (E-value= 9e-98), and contains all domains expected for a bacterial primase (**Figure S5**). We purified the *C. difficile* primase without an affinity tag to > 95% purity. SEC-MALS/dRI analysis showed a single peak corresponding to a molecular weight (MW) of 74kDa, indicating that the primase is monomeric under the conditions tested (expected MW 70 kDa) (**Figure S6**).

### C. difficile *helicase is activated by primase*

We performed helicase assays to determine the effect of primase on helicase acitivity.

First, we determined if the CD3657 helicase displayed activity in this particular assay in the absence of any other proteins. Approximately 20% of the oligonucleotide was displaced after as little as 2 minutes of incubation, and this fraction remained stable over the course of 35 minutes (Figure 4A). This suggests that helicase may be capable of self-loading but displays marginal DNA unwinding activity by itself.

**Figure 4.**
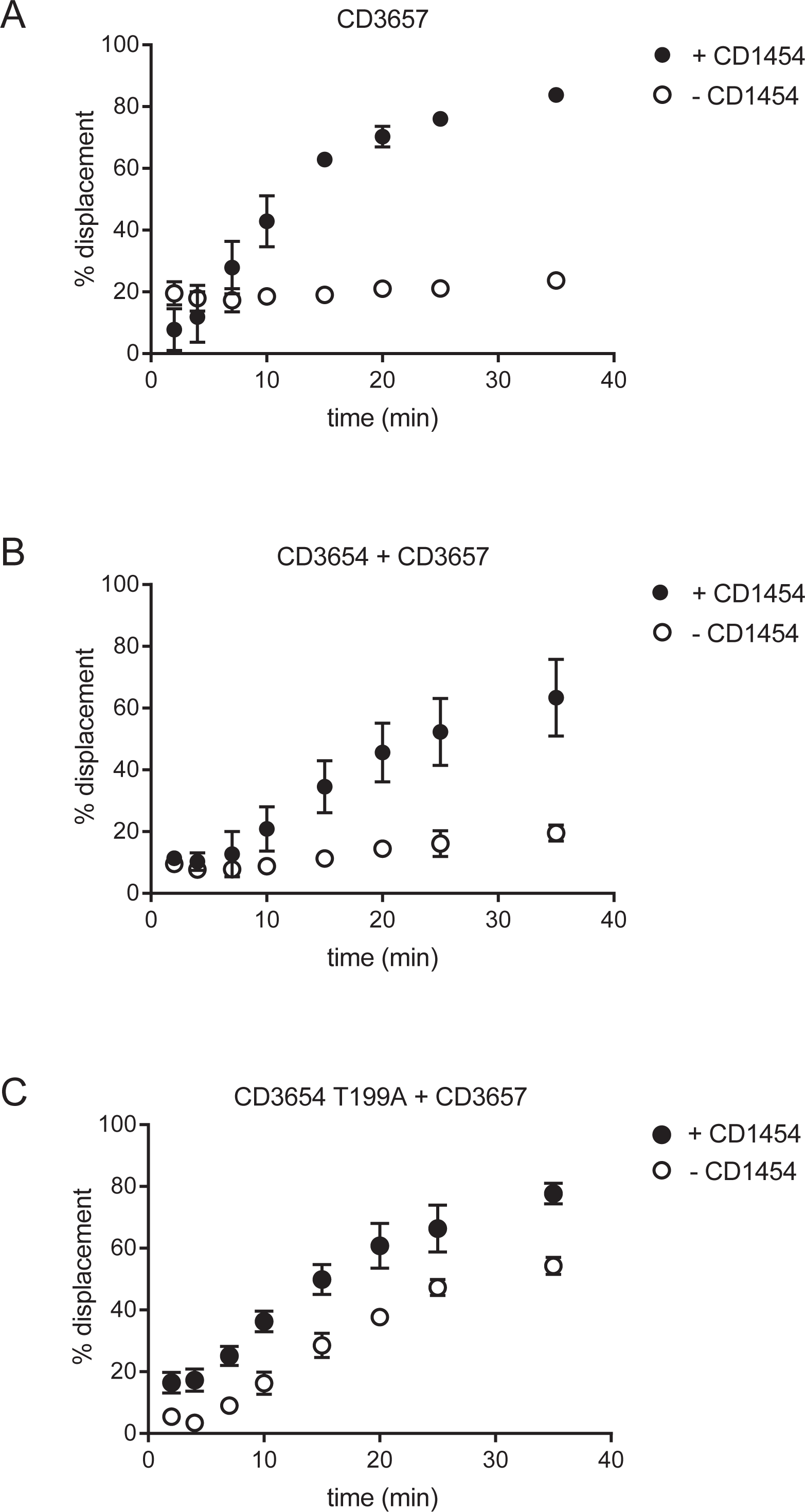
Helicase activity is stimulated by primase. Helicase activity was assayed by quantifying the displacement of a radiolabelled (γ^32^P-ATP) oligonucleotide partially annealed to the single stranded circular DNA m13mp18. Percent displaced signal from the helicase assays in time. **A.** Helicase activity of the CD3657 helicase with and without the CD1454 primase. **B.** Helicase activity of the CD3657 helicase in the presence of the putative loader protein CD3654, in the presence and absence of the CD1454 primase. **C.**Helicase activity of the CD3657 helicase in the presence of a representative Walker A mutant (T199A) of the putative loader protein CD3654, in the presence and absence of the CD1454 primase. The other mutants of CD3654 (K198R, D258Q) gave similar results (**Supplemental Figure 7**) but have been omitted for clarity.

When the CD1454 primase and CD3657 helicase were combined in equimolar amounts, a 3.5 fold increase in displacement of the oligonucleotide (up to ~80%) was observed after 35 minutes (Figure 4A). This indicates that primase has a profound stimulatory effect on helicase activity in this assay. Strikingly, strand displacement by helicase seems inhibited in the presence of primase compared to helicase alone at early time points (<10 minutes). This suggests that primase may inhibit self-loading, in addition to its role as an activator of helicase activity. Control experiments using only primase did not result in significant displacement of the oligonucleotide, demonstrating that the displacement is not the result of an inherent property of the primase protein (unpublished observations).

The results from the helicase assay suggest a functional interaction between the helicase and primase proteins. We tried to validate the interaction using a bacterial two-hybrid system. Despite of our efforts, no interaction could be demonstrated between the full-length primase and helicase proteins, or the helicase interacting domain of primase and helicase in bacterial two hybrid experiments (unpublished observations). This suggests that the interaction between primase and helicase may be very weak (below detection limit of the assay) and/or transient, concordant with observations in *E. coli* (46), or that the bacterial two-hybrid system does not allow recapitulation of the conditions necessary for the interaction between the helicase and primase proteins.

### The putative loader protein CD3654 "locks" the CD3657 helicase

Next, we set out to investigate the role of the putative loader CD3654 on the helicase activity of CD3657 in the presence and absence of primase. We observed a very low level of displacement (<20%) in our helicase assay in the presence of both helicase and the putative loader (Figure 4B). Notably, within the first 10 minutes of the assay the fraction of displaced oligonucleotide did not exceed 10%, in contrast to the situation with helicase alone where it reached 20% (Figure 4A). At end point, the fraction was comparable between the two conditions. This suggests that – at least at early time points – the presence of the putative loader negatively affects helicase activity.

We wondered if the positive effect of the CD1454 primase on CD3657 helicase activity could be observed in the presence of both the putative loader ATPase CD3654 and helicase. From 8 minutes on, a clear stimulation of helicase activity by primase was observed, reaching ~60% displacement over a time course of 35 minutes (Figure 4B). Interestingly, the addition of the putative loader protein resulted in a 20 percent reduction in strand displacement over a time-course of 35 minutes compared to results obtained with a combination of helicase and primase (~80%) (Figure 4A). We conclude that primase can activate helicase activity in the presence of the putative loader, but that the loader retains a negative effect on overall helicase activity in this assay.

To exclude the possibility that the displacement observed in our helicase assays could be attributed to another protein than the CD3657 helicase, we used the CD3657 K214R and T215A mutants. We observed only background levels (<5%) of displacement in the presence of CD3657 mutant helicases, primase and the putative loader (**Figure S7**) in comparison to ~60% strand displacement for the wild type helicase in the presence of the same proteins (Figure 4B).This shows that the displacement observed in our experiments can be attributed to helicase alone and not some other factor.

In *E. coli*, interaction between the loader ATPase protein and helicase does not require ATP binding by the loader (8;70;71). However, hydrolysis of ATP to ADP by the loader (that results in dissociation from the helicase-loader complex) appears crucial to lift the negative effect of the loader on helicase activity (8;70;71). Therefore, we set out to investigate the effect of CD3654 proteins with a mutated Walker A or Walker B motif on helicase activity. The lysine residue in (K198) in the Walker A motif is predicted to directly interact with ATP, and mutation of homologous residues is known to eliminate appropriate ATP binding of AAA+ proteins, resulting in inactivation (Hanson 2005). The threonine residue in Walker A (T199) and the aspartic acid residue in Walker B (D258Q) are predicted to be involved in coordination of an Mg^2+^ ion within the ATP-binding site. Overall, the three mutants (K198R, T199A and D258Q) should be affected in their ability to coordinate and/or hydrolyze ATP.

We found a 2.5-fold higher displacement (~50%) of the oligonucleotide in the helicase assays with all three CD3654 Walker A and B mutants **(Figure 4C and Figure S8**) compared to the wild type (Figure 4B) in the absence of primase. Interestingly, helicase activity in assays combining wild type CD3657 helicase, CD1454 primase, and mutants of the putative loader was similar (~80%, Figure 4C) to the activity measured in the two protein helicase-primase experiment (Figure 4A). This suggest that binding and/or hydrolysis of ATP by the putative loader is not required to deliver the CD3657 helicase to the DNA, but is at least partially responsible for the negative effect (“locking”) of the helicase activity.

### C. difficile *primase trinucleotide specificity is similar to* A. aeolicus *primase*

Above, we have established a crucial role for the *C. difficile* primase CD1454 in the activation of the CD3657 helicase. Next, we sought to evaluate primase activity. To confirm enzymatic activity and identify the template initiation motif, two 50-mer oligonucleotides that comprised all 64 potential trinucleotide sequences were used in a priming assay, as previously described (72). These results confirmed enzymatic activity and revealed likely trinucleotide motifs that supported *de novo* primer synthesis (unpublished observations). Next, a more detailed analysis of template specificity was performed using a 23-mer ssDNA-template containing the trinucleotide of interest (Figure 5A). We found that the CD1454 primase preferentially initiated *de novo* RNA primer synthesis on the initial 5’-d(CCC) motif within the 23-mer ssDNA template, producing a 17-mer rather than an anticipated 16-mer RNA primer (Figure 5B). As a negative control, the 5’-d(ACA)-specific 23-mer ssDNA was tested in which no priming by *C. difficile* primase occurred (Figure 5B). The activity of the CD1454 was inhibited by highMg^2+^ concentrations (>20 mM), and largely independent of NTP concentration(>1 mM) under the conditions of the assay (**Figure S9**).

**Figure 5.**
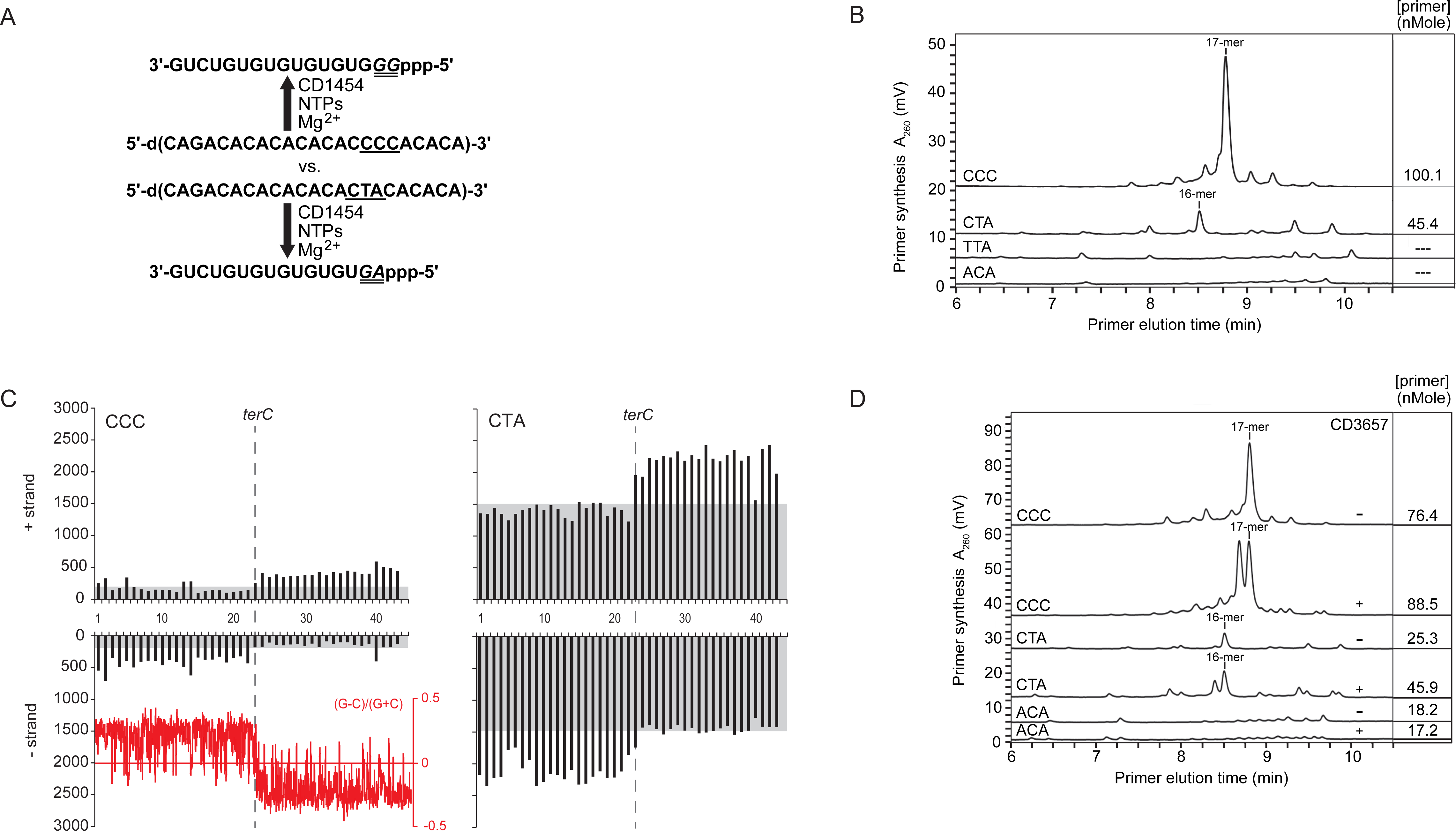
CD3657 affects trinucleotide specificity of CD1454. **A**. Schematic depiction of the thermally denaturing high-performance liquid chromatography analysis of primase activity. **B**. Analysis of CD1454 priming activity at different trinucleotides. **C**. The number of CCC and CTA trinucleotides on the + and – strand of the *C. difficile* 630Δ*erm* was calculated in 100000 bp bins. Skew in trinucleotide frequency is highlighted using a gray box. The position of the putative terminus of replication (*terC*) is indicated with a vertical dashed line. The red inset shows the GC skew, as calculated in Artemis (99) with a window size of 5000. **D**. CD3657 affects CD1454 primase activity. The amount of primer produced by CD1454 in the presence (+) or absence (-) of CD3657 on ssDNA templates with different trinucleotides at stoichiometric concentrations of [CD3657]_6_:[CD1454]_3_.

The observed template specificity for the CD1454 primase differs from other bacterial primases. More specifically, 5’-d(CTA) only supported minimal priming and no initiation occurred on 5’-d(TTA) by the *C. difficile* primase (Figure 5B), whereas substantial dinucleotide polymerization occurred on these two trinucleotides by primases from other Firmicutes such as *G. stearothermophilus*, *S. aureus*, and *Bacillus anthracis* (43;52;73). The trinucleotide 5’-d(CTG) was also not an effective template for *C. difficile* primase (unpublished observations), whereas it is efficiently primed by primase from Gammaproteobacteria such as *E. coli*, *Pseudomonas aeruginosa*, and *Yersinia pestis* (73-75). To date, only the primase from the hyperthermophile *Aquifex aeolicus* has been shown to initiate primer synthesis on the trinucleotide 5’-d(CCC) (72).

Since primase presumably scans the ssDNA template in the 3’ to 5’ direction until an initiation trinucleotide is encountered, the influence of the 5’ nucleotide adjacent to the preferred 5’-d(CCC) trinucleotide motif was evaluated. We found that CD1454 primase was most efficient at initiating *de novo* primer synthesis when the 5’ nucleotide was a cytosine or - with a slightly lower efficiency - a thymine (**Figure S10**). Considerably less priming occurred when the 5’ adjacent nucleotide contained an adenine base and only marginal nonspecific priming occurred when this nucleotide contained a guanine (**Figure S10**). These results suggest that the nucleotide 5’ to the preferred trinucleotide influences the catalytic activity at the active site for subsequent initiation of *de novo* primer synthesis.

### Helicase influences trinucleotide specificity of primase

The identification of the 5’d(CCC) motif as the preferred trinucleotide for initiation by the CD1454 primase was unexpected, as the *C. difficile* chromosome has a G+C content of only 29% (64) and other low-GC Firmicutes, such as *S. aureus* and *G. stearothermophilus* (43;52) preferentially initiate at 5’d(CTA). Therefore, we evaluated the relative frequency of the CCC motif on the plus and minus strand of the *C. difficile* chromosome and compared it to the relative frequency of CTA motif on which the CD1454 initiates less efficiently (Figure 5C). Our analysis showed that CTA trinucleotides were on average five to ten-fold more frequent within the *C. difficile* chromosome than the preferred CCC-motif. Strikingly, there appears to be a strand bias in the occurrence of the motifs, that mirrors the GC-skew ([G-C]/[G+C]) of the chromosome (Figure 5C). This suggests that the motifs are preferentially associated with the lagging strand where primase acts and indicate a possible role for primase in generating the strand bias.

As it has previously been shown that helicase can stimulate primase activity, affect primer length and modulate trinucleotide specificity in *E. coli* (47;48;50), we determined whether the CD3657 helicase could enhance priming at the non-preferred trinucleotide. The effect of CD3657 helicase on CD1454 primase activity was evaluated using 23-mer ssDNA templates that either contained the preferred trinucleotide 5’-d(CCC), the Firmicute-preferred trinucleotide 5’-d(CTA), or 5’-d(ACA) (negative control). RNA primer production by the primase CD1454 was strongly stimulated on 5’-d(CTA) in the presence of the CD3657 helicase (45.9 vs 25.3 nMole) at stoichiometric concentrations of [CD3657]_6_:[CD1454]_3_. Helicase-stimulated primase activity on 5’-d(CCC) increased RNA primer synthesis by only 1.15-fold and, as expected, no stimulation of RNA primer synthesis occurred on the 5’-d(ACA) trinucleotide (Figure 5D). We also observed that helicase stimulated primase to synthesize slightly shorter RNA polymers (Figure 5D). The production of a shorter primer compared to a longer primer by helicase-stimulated primase would allow for a faster rate of extension by DNA polymerase while still providing the required free 3’ hydroxyl group for initiation, as occurs *in vivo*. Collectively, our data show that the stimulatory effect of helicase on primase activity *in vitro* is clearly enhanced on the trinucleotide 5’-d(CTA), which is preferred by other Firmicute primases studied to date, whereas the effect at the *C. difficile*-preferred 5’-d(CCC) sequence was only minimal, most likely due to the already high efficiency of priming on this motif by the CD1454 primase.

We conclude that through the interaction of primase with helicase and the relative abundance of the 5’d(CTA) the overall efficiency of priming is likely greatly enhanced.

### A lysine residue contributes to trinucleotide specificity of primase

Considering that the *C. difficile* primase trinucleotide specificity (Figure 5B) is unusual for Firmicutes, but resembles that of the hyperthermophilic *A. aeolicus* primase (Larson 2008), we wanted to determine the factors that might contribute to this trinucleotide template specificity for subsequent dinucleotide polymerization.

We performed pair-wise sequence comparisons between the zinc binding domain (ZBD) that is involved in sequence specific DNA binding of primase in *C. difficile* and ten other bacterial species (*Clostridium perfringens, G. stearothermophilus*, *S. aureus*, *B. anthracis*, *B. subtilis*, *E. coli*, *P. aeruginosa*, *Y. pestis*, *A. aeolicus*, and *Francisella tularensis*). *C. perfringens* (53.8% identity and 87.9% similarity in 91 residues) had the highest amino acid sequence homology followed by *B. anthracis* (49.5% identity and 87.9% similarity in 91 residues) and *A. aeolicus* (47.2% identity and 83.1% similarity in 89 amino acids). The *E. coli* ZBD had the lowest homology (43.2% identity and 83.0% similarity in 88 residues). The Clustal Omega alignment tool (http://www.ebi.ac.uk/Tools/msa/clustalo) placed the *C. difficile* CD1454 ZBD closest to the ZBD of *A. aeolicus* and *G. stearothermophilus* (Figure 6A). The (partial) clustering of the *C. difficile* CD1454 primase with other bacterial primases that have different trinucleotide specificity led us to anticipate that the composition and spatial location of specific amino acids in the ZBD relative to the RNA polymerase domain in primase must be critical for template recognition and phosphodiester bond formation.

**Figure 6.**
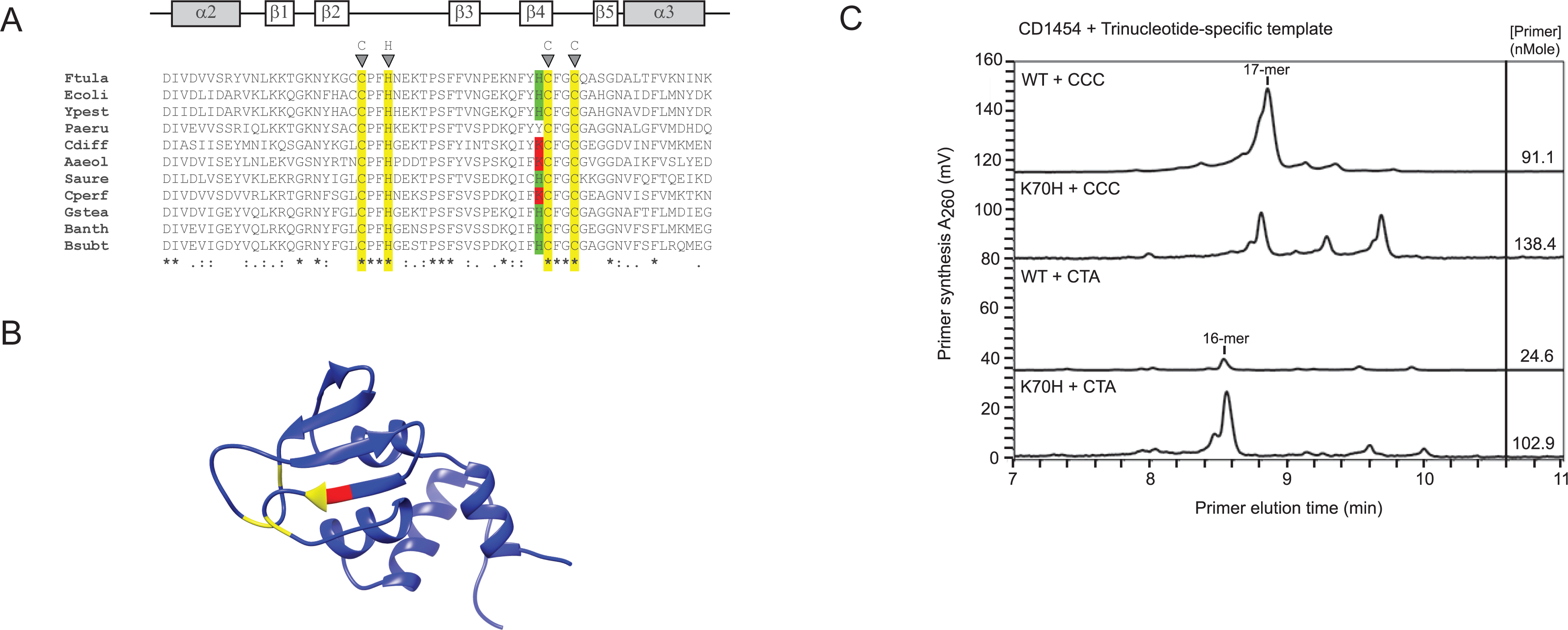
A lysine residue affect trinucleotide specificity of CD1454. **A.** Alignment of the α2-α3 region of the zinc binding domain of primases with characterized trinucleotide specificity. Predicted secondary structure (alpha helix in gray, beta sheet in white) is indicated above the alignment. Ftula: *Francisella tularensis*, Ecoli: *Escherichia coli*, Ypest: *Yersinia pestis*, Paeru: *Pseudomonas aeruginosa*, Cdiff: *Clostridium difficile*, Aaeol: *Aquifex aeolicus*, Saure: *Staphylococcus aureus*, Cperf: *Clostridium perfringens*, Gstea: *Geobacillus stearothermophilus*, Banth: *Bacillus anthracis*, Bsubt: *Bacillus subtilis*. Highlighted are the zinc coordinating residues of the CHC2 zinc binding motif (yellow), and the adjacent histidine (green) or lysine (red) residues. Sequence conservation is indicated below the alignment. **C**. Rotated view of the CD1454 primase zinc binding domain in **A** indicating the zinc coordinating residues (yellow) and the location of the lysine residue (red). **C**. Thermally denaturing high-performance liquid chromatography analysis of primase activity of wild type (WT) and K70H mutant CD1454 primase protein. The mutation affects RNA primer synthesis from both preferred 5’-d(CCC) and non-preferred 5’-d(CTA) trinucleotides but shows the strongest increase in activity from the 5’-d(CTA) trinucleotide.

In the multiple sequence alignment, a lysine (K) at residue 70 in the ZBD of *C. difficile* primase was unique to *C. difficile*, *C. perfringens*, and *A. aeolicus* (Figure 6A). Structural modelling of CD1454 with Phyre2 (76) revealed that this particular lysine residue is in close proximity of the zinc ribbon motif that tetrahedrally coordinates a zinc ion and is essential for primase function (**Figure S5 and 6B**) (77). Strikingly, the PDB template used by Phyre2 is 2AU3, the *A. aeolicus* primase (53). The crystal structure of *G. stearothermophilus* DnaG primase contains a histidine at this position (49;77) and the multiple sequence alignment suggests that other Firmicutes do as well. We hypothesized that the exposed lysine residue influences primase initiation specificity. To address this experimentally, we mutated the K70 residue of the CD1454 primase to a histidine. If the lysine contributes to the unusual trinucleotide specificity of the *C. difficile* primase, we would expect reduced priming on the 5’-d(CCC) motif by the CD1454 K70H mutant compared to the wild type CD1454 primase. Indeed, CD1454 K70H showed substantially reduced initiation on the 5’-d(CCC) motif and enhanced priming on the 5’d(CTA) motif (Figure 6C). The CD1454 K70H synthesized slightly more primers that wild type primase due to relaxed template specificity, as evidenced by the synthesis of RNA polymers longer than 17 nucleotides in length (Figure 6C).

Importantly, these results demonstrate that modifying a single residue within the ZBD of the CD1454 primase is sufficient to alter initiation specificity and suggests that the exposed lysine residue is crucial for preferential initiation on the 5’-d(CCC) motif.

## DISCUSSION

*In silico* analysis of the *C. difficile* genome by BLASTP identified homologs of most proteins that are involved in the DNA replication process in *B. subtilis*, which is generally considered the model for Gram-positive bacteria. However, *C. difficile* does not encode a BsDnaB homolog. This protein, together with BsDnaD and BsDnaI, is strictly required for helicase loading in *B. subtilis in vivo* (11-13). A weak homologue of DnaD was identified that may be involved in DNA replication (CD3653; e-value 4e-05), although query coverage (47%) and identity (29%) were low. This situation is reminiscent of that in some *Mollicutes*, where analyses also have identified only a *DnaD*-like gene (14). Despite a lack of clear homology at the primary amino acid sequence level, DnaB and DnaD are structural homologs (17). Fusions of these proteins are found in phage-related replication proteins and it was suggested that in the absence of DnaB, a single fusion protein may couple or combine both functions (14;65). Nevertheless, the situation in *C. difficile* differs from those in phage and *Mollicutes*. Structure predictions reveal that the phage-related and the *Mollicute* DnaD-like proteins have a two-domain structure containing one copy of the DDBH1 and DDBH2 domain (17). Moreover, the proposed hybrid function of phage proteins is based on limited local amino acid sequence similarity in the C-terminal (DDBH2) domain only (65). CD3653 on the other hand has a three domain structure with a single DDBH1 and two DDBH2 domains, like DnaB, despite the lack of clear sequence similarity to this protein (Figure 1). It is tempting to speculate that CD3653 in *C. difficile* may perform functions similar to both DnaD and DnaB in *B. subtilis*. DDBH2 domains are characterized by an YxxxIxxxW motif (17). In BsDnaB, this motif is degenerate in the first DDBH2 domain. In contrast, the motif is readily identified in both DDBH2 domains of CD3653 (Figure 1). Our data are consistent with a model where an ancestral three-domain DnaD-like protein was duplicated and subsequently diverged in certain Firmicutes like *B. subtilis*.

DnaB-like helicases (note that this nomenclature is based on the *E.coli* protein name) belong to the superfamily IV of DNA helicases, and the functional unit of this protein is a hexamer (25-27;33). In *E. coli*, the helicase is found to be a stable hexamer over a broad protein concentration range of 0.1 to 10 µM (78) and is active as a pre-formed multimer. Helicases belonging to the ring-maker class, such as BsDnaC of *B. subtilis*, can occur in a low-oligomeric or monomeric state under dilute conditions (5;13). Our experiments indicated that helicase is monomeric in the low-micromolar or nanomolar range (**Figure S1**), which is likely reflective of the intracellular concentration of protein. *C. difficile* CD3657 can form hexameric assemblies at higher concentrations (**Figures S1 and S2**), but these pre-formed hexamers are inactive, even in the presence of primase (our unpublished observations), in contrast to the situation in *E. coli*. Our data are therefore consistent with the notion that CD3657 belongs to the ring-maker class of helicases (5).

Although the addition of ATP was not strictly required for hexamerization of CD3657 at high protein concentrations (**Figure S2**), the interaction between putative helicase and loader protein was found to be ATP-dependent (Figure 2). Mutations in the conserved Walker A and B motifs of the putative loader protein did not fully abrogate the interaction with the wild type helicase (**Figure S3**). Similarly, in *E. coli*, nucleotide binding to the helicase loader was not a pre-requisite for association with helicase (8;70;79). Instead, our data indicate that the association of ATP with helicase is crucial for the interaction with the loader protein (Figure 3). Notably, there is a correlation between the ability of the helicase to interact with the loader protein and form homohexamers, since a T215A (Walker A) mutant of helicase is defective for both, at least under dilute concentrations of helicase (Figure 3). In contrast, the equivalent mutation in *G. stearothermophilus* helicase (T217A) does not affect its ability to form hexamers (58), and the interaction of this protein with *B. subtilis* DnaI readily occurs in the absence of ATP (80).

Both Walker A mutants of CD3657 demonstrate similar effects on the protein-protein interactions, that we attribute to defects in ATP binding rather than hydrolysis (Figure 3). We find that a Walker B mutant (CD3657 D318A), but not a mutant that is predicted to be able to bind but not hydrolyse ATP (CD3657 E239A) mirrors our findings with the Walker A mutants, at least in a bacterial two hybrid assay (unpublished observations). CD3657 T215 is predicted to be involved in Mg^2+^ binding, which is necessary for stabilizing ATP binding. Binding of the nucleotide to helicase is associated with conformational changes; the N-terminal collar domain constricts upon nucleotide binding in *A. aeolicus* and to a lesser extent *E. coli.* This constricted conformation is believed to favor an interaction with the loader protein (81) and it is therefore conceivable that in the absence of ATP, the *C. difficile* helicase adopts a (dilated) conformation that is incompatible with a functional interaction with the putative loader protein.

Despite substantial bioinformatic and biochemical evidence that CD3654 is indeed the *C. difficile* helicase loader, we found an inhibitory effect of the CD3654 on the activity of the CD3657 *in vitro*, at least in the presence of the CD1454 (Figure 4). At first, the negative effect of CD3654 seems at odds with its proposed role as loader. However, in *E. coli* it has been demonstrated that ATP-bound EcDnaC loader protein can act as an inhibitor of the EcDnaB helicase (8;70;71). In our experiments we find that mutants of CD3654 appear capable of loading and/or activating the helicase, but are defective in restraining the DNA-unwinding activity of the helicase like wild type CD3654 (Figure 4C and **Figure S8**). Moreover, a functional role for CD3654 in the essential process of helicase loading and/or activation is supported by the observation that no transposon insertions were obtained in the homolog of the *cd3654* in an epidemic strain of *C. difficile* (R20291_3513) (82).

Our data strongly suggest that the presence of loader protein alone is not sufficient to load and activate the helicase of *C. difficile*, and that at least one other factor is needed to reconstitute its activity. Indeed, we determined that primase is required as an activator of helicase activity in the presence of the loader (Figure 4).

Helicases are complex proteins, and their properties can both alter and be altered by other replication factors (81). DnaB-like helicases consist of two-tiered homo-hexameric rings, one assembled from six subunits of the C-terminal domain and the other formed by the N-terminal domains. The helicase loader interacts with the C-terminal ATPase domain (61;70;83), and the same domain is required for the interaction with the τ subunit of the clamp loader in *E.coli* (84), and *B. subtilis* (85;86). Strikingly, for the latter it has been shown that the interaction depends on ATP binding, as a BsDnaC T217A mutant fails to form a complex (86). This finding is very similar to our observations for the interaction between the *C. difficile* helicase and loader. The N-terminal domain of helicase forms a platform for the interaction with primase in both *E.coli* and *B. stearothermophilus* (61;81). Unlike the helicase loader, binding of primase to helicase is promoted by a dilated conformation of the N-terminal domain that exposes the interaction surface (57;81). The helicase-primase interaction is mutually stimulatory, with distinct but overlapping networks of residues in helicase responsible for the modulation of either helicase or primase activity (43;87;88). Primase binding counteracts the binding of the loader protein in *E. coli* (60). Similarly, the helicase loader from *B. subtilis* was found to dissociate from the complex when primase and polymerase bind to helicase in gel-filtration experiments (Rannou 2013). We find that CD1454 stimulates CD3657 helicase activity (Figure 4).

Loading of the CD3657 helicase differs from the situation in other Gram-positive bacteria. Most notably, CD3657 seems to be capable of self-loading, has increased helicase activity in the presence of primase (CD1454) and is negatively influenced by the presence of the putative loader protein (Figure 2A). *B. stearothermophilus* helicase demonstrates significant helicase activity by itself (58), and the *B. subtilis* helicase is strongly activated by its cognate loader (16;56). Instead, helicase loading in *C. difficile* is somewhat reminiscent of the situation in the Gram-negative *H. pylori*, where a dodecameric self-loading helicase remains inactive until activated by primase, leading to the dissociation of the dodecamer into two hexamers (15). However, we did not find any evidence for dodecameric assemblies for CD3657.

Helicase loading in *C. difficile* also seems to differ in crucial aspects from the Gram-negative model *E. coli*. Walker A mutants of the EcDnaC loader protein are capable of loading the EcDnaB helicase but do not sustain helicase activity, suggesting that ATP turnover by the loader is required to release the helicase (8). Our data show that in *C. difficile*, (self) loading of CD3657 is stimulated by Walker A mutants of CD3654, but that these mutants of the loader readily release active helicase (Figure 2C). This suggests a role for ATP binding and/or hydrolysis in “locking” helicase activity.

Moreover, EcDnaB helicase is in complex with the EcDnaC loader protein throughout the loading process and remains inactive until the EcDnaG primase binds to helicase, thereby releasing the loader protein (60). Our data do not exclude a similar role for primase in helicase loading of *C. difficile*, but do show that the role of the CD1454 primase is not limited to the release of the putative loader; very strong stimulation of CD3657 helicase activity is observed when CD1454 primase is added in the helicase assay in the absence of the CD3654 loader (Figure 4A). The CD1454 primase may still stimulate ATP turnover by the CD3654 loader protein. We consider two possible, not mutually exclusive, scenarios to explain the activation of helicase by primase in the absence of the loader. First, primase may stabilize the hexameric helicase on the DNA, which indirectly contributes to the unwinding activity. Second, primase may act as a direct activator of the DNA-unwinding activity of helicase. Our experiments do not discriminate between these possibilities, though stabilization of the hexamer or other conformational changes in the hexameric helicase induced by primase have also been observed in *E. coli* (40;57;59;60).

We found that the primase of *C. difficile* has an unusual trinucleotide specificity, with a preference for 5’-d(CCC) (Figure 5), similar to *A. aeolicus* (72). This is in contrast with primases from other Firmicutes, which initiate *de novo* primer synthesis on 5’-d(CTA) and 5’-d(TTA), and Gammaproteobacterial primases, which initiate on 5’-d(CTG) and 5’-d(CTA)(49). The first two nucleotides in all of these preferred and recognized trinucleotides are pyrimidines (C or T). As they serve as the template for the corresponding dinucleotide, the first two nucleotides in the RNA primer will be purines (G or A). The levels of both ATP and GTP directly or indirectly provide a means by which bacteria can sense the energy status of the cell (89-92), and the nucleotide preference might ensure a coupling between the efficiency of lagging strand DNA synthesis and nutritional status of the cell.

Priming by the primase CD1454 was highest when the nucleotide 5’ adjacent to the preferred 5’-d(CCC) trinucleotide was a pyrimidine (**Figure S10**), consistent with a previously hypothesized context dependent enzyme activity (72). Our results indicate that pyrimidines, probably in part due to the smaller size compared to purines, provide the optimal context for catalysis and dinucleotide polymerization.

We probed the origin of primase trinucleotide specificity and found that a lysine (K) adjacent to the zinc-ribbon motif of CD1454 is important for the optimal physico-chemical environment for primer initiation on 5’-d(CCC) (Figure 6). Based on this finding, we would predict a similar specificity for the primase of *Clostridium perfringens*. Unfortunately, we could not confirm this result with *C. perfringens* primase so far. We purified the CPF_2265 primase from *C. perfringens* ATCC133124 but our attempts to obtain a primase with priming activity have failed thus far.

A 5’-d(CCC) trinucleotide specificity has previously only been observed for *A. aeolicus* (72). Despite the predicted structural homology between the *A. aeolicus* and *C. difficile* primases (Figure 6B), this was unexpected, as *C. difficile* is a mesophilic, spore-forming, Gram-positive pathogen, whereas *A. aeolicus* is a hyperthermophilic Gram-negative bacterium. However, cladistic studies using multiple signature proteins indicate that the *Aquifex* lineage emerged from Gram-positive bacteria, prior to the split of Gram-positive and Gram-negative bacteria (93;94). These studies concluded that the Firmicutes are presumably among the most ancient bacteria and that the Aquificales have diverged much later in evolution (93). Indeed, an analysis of the GC-content of rRNA clusters suggests that hyperthermophilic species have evolved from mesophilic organisms via adaptation to high temperature (95) and that the Gram-negative double membrane may have been derived from sporulating Gram-positives (96;97). Thus, the study of *C. difficile* may also contribute to a better understanding of the evolution of the bacterial DNA replication machinery and facilitate the development of antibiotics that target these essential proteins in pathogenic bacteria.

## ACKNOWLEDGEMENT

Plasmids pKEK1286 and pKEK1287 were kindly provided by K.E. Klose (University of Texas at San Antonio). D. Jex and J. Gibson are acknowledged for exploratory helicase activity assays and analytical gel filtrations.

We thank J.B. van Voorden for performing the site-directed mutagenesis on our constructs, R. Scherrers from Wyatt Technology for the SEC-MALS analysis of the CD1454 protein, and D. Tomkiewicz for help with preparing Figure S6.

## Supplemental Information

A short description of available Supplemental Information is given below.

- Supplementary Methods.
- Figure S1. CD3657 demonstrates concentration dependent hexamerisation.
- Figure S2. ATP dependent interaction of the helicase CD3657 and the putative loader CD3654 at high concentrations of proteins.
- Figure S3. Walker A mutants of CD3654 retain the ability to interact with CD3657.
- Figure S4. Helicase activity in the presence of helicase and (putative) loader proteins of *B. subtilis* and *C. difficile*.
- Figure S5. Phyre2 model of the CD1454 primase.
- Figure S6. Molecular mass determination of the CD1454 primase protein.
- Figure S7. Helicase activity is abrogated in a Walker A mutant of CD3657.
- Figure S8. Helicase activity is not inhibited in the presence of mutant CD3654 loader proteins.
- Figure S9. Effect of Mg2+ and NTP concentration on CD1454 activity.
- Figure S10. The effect of the 5’ flanking base on priming efficiency of CD1454 at the preferred trinucleotide.
- Table S1. Oligonucleotides used in this study.
- Table S2. Plasmids used in this study.

